# Intrinsic Dynamics and Neural Implementation of a Hypothalamic Line Attractor Encoding an Internal Behavioral State

**DOI:** 10.1101/2024.05.21.595051

**Authors:** Amit Vinograd, Aditya Nair, Scott W. Linderman, David J. Anderson

## Abstract

Line attractors are emergent population dynamics hypothesized to encode continuous variables such as head direction and internal states. In mammals, direct evidence of neural implementation of a line attractor has been hindered by the challenge of targeting perturbations to specific neurons within contributing ensembles. Estrogen receptor type 1 (Esr1)-expressing neurons in the ventrolateral subdivision of the ventromedial hypothalamus (VMHvl) show line attractor dynamics in male mice during fighting. We hypothesized that these dynamics may encode continuous variation in the intensity of an internal aggressive state. Here, we report that these neurons also show line attractor dynamics in head-fixed mice observing aggression. We exploit this finding to identify and perturb line attractor-contributing neurons using 2-photon calcium imaging and holographic optogenetic perturbations. On-manifold perturbations demonstrate that integration and persistent activity are intrinsic properties of these neurons which drive the system along the line attractor, while transient off-manifold perturbations reveal rapid relaxation back into the attractor. Furthermore, stimulation and imaging reveal selective functional connectivity among attractor-contributing neurons. Intriguingly, individual differences among mice in line attractor stability were correlated with the degree of functional connectivity among contributing neurons. Mechanistic modelling indicates that dense subnetwork connectivity and slow neurotransmission are required to explain our empirical findings. Our work bridges circuit and manifold paradigms, shedding light on the intrinsic and operational dynamics of a behaviorally relevant mammalian line attractor.

## Introduction

Neural computations have long been studied from two distinct vantage points. One focuses on understanding behaviorally specialized neuron types and their functional connectivity^1–3^, while the other investigates emergent properties of neural networks, such as attractors^4–6^. Attractors of different topologies are theorized to encode a variety of continuous variables, ranging from head direction^7^, location in space^8^ and internal states^9^. Recent data-driven methodologies have allowed for the discovery of such attractor mediated computations directly in neural data^9–12^. Consequently, attractor dynamics have received increasing attention as a major type of neural coding mechanism^7,8,13 4,10^.

Despite this progress, establishing that these attractors arise from the intrinsic dynamics of the observed network remains a formidable challenge^4,8^. Unaccounted external inputs such as feedforward synaptic input can profoundly influence computational dynamics observed at a given site^13^. Therefore, experimental perturbations are pivotal to determine whether observed attractor dynamics are locally computed or inherited. This calls for combining large-scale recordings with perturbations of neuronal activity *in vivo*. While this has been accomplished for a point attractor that controls motor planning in cortical area ALM^14,15^, spatial ensembles that regulate short term memory^16,17^, and for a ring attractor in *Drosophila*^18,19^, there is no study reporting such perturbations for a continuous attractor in any mammalian system. While theoretical work on continuous attractors in mammals is well-developed^8^, the lack of direct, neural perturbation-based experimental evidence of such attractor dynamics has hindered progress towards a mechanistic circuit-level understanding of such emergent manifold-level network features^4^.

VMHvl^Esr1^ neurons comprise a key node in the hypothalamic-extended amygdala social behavior network and have been causally implicated in aggression^20,21^. Calcium imaging of these neurons in freely behaving animals has revealed mixed selectivity, with aggression sparsely represented at the single-neuron level^22,23^. Yet aggressive behavior can be accurately decoded from population activity^23^, raising the question of which aspect(s) of this activity contain such information. Application of dynamical system modeling has revealed an approximate line attractor in VMHvl that correlates with the intensity of agonistic behavior, suggesting a population-level encoding of a continuously varying aggressive internal state^9^. This raises the question of whether the observation of a line attractor in a dynamical systems model fit to VMHvl^Esr1^ neuronal activity reflects intrinsic dynamics, or rather passive inheritance of such dynamics from an upstream source.

This question can be addressed, in principle, using all-optical methods to observe and perturb line attractor-relevant neural activity^4,24–26^. A challenge in applying these methods during aggression is that current technology requires head-fixed preparations, and head-fixed mice cannot fight. To overcome this challenge, we exploited the recent observation that VMHvl^PR^ neurons (which overlap Esr1 neurons) mirror inter-individual aggression.^27^ Here we show that VMHvl^Esr1^ neurons also exhibit line attractor dynamics during the passive observation of aggression, and that such neurons are largely overlapping with line attractor-contributing neurons in attacking mice. Leveraging this mirror paradigm to generate line attractor dynamics in head-fixed subjects, we performed dynamical model-guided, closed-loop perturbations of VMHvl^Esr1^ activity. This approach revealed that the VMHvl line attractor indeed reflects intrinsic neural dynamics in this nucleus. Furthermore, it identified a neural implementation rooted in selective functional connectivity within attractor-weighted ensembles that is likely mediated by slow neurotransmission, ensuring the attractor’s stability. Collectively, our findings elucidate, for the first time, a circuit-level foundation for a continuous attractor in the mammalian brain.

## Results

### Line attractor dynamics during observation of aggression

Recent studies have demonstrated that VMHvl contains neurons that are active during passive observation of as well as active participation in aggression, and that re-activating the former can evoke aggressive behavior^27^. However, those findings were based on a very small number of VMHvl^PR^ neurons, which might comprise a specific subset distinct from those contributing to the line attractor (the latter represent ∼20-25% of Esr1^+^ neurons^9^). To assess whether these mirror-like responses can be observed in those Esr1^+^ neurons that contribute to line attractor dynamics, we performed microendoscopic imaging^28^ of VMHvl^Esr1^ neurons expressing jGCaMP7s in the same freely moving animals sequentially during engagement in, and observation of, aggression (Extended Data 1a-e). Analysis using recurrent switching linear dynamical systems (rSLDS)^29^ to fit a model to each dataset revealed an approximate line attractor under both conditions, exhibiting ramping and persistent activity aligned and maintained across both performed and observed attack bouts (Extended Data 1g-p). Notably, the integration dimension aligned with the line attractor (“x_1_”) was weighted by a consistent set of neurons under both conditions, suggesting that a highly overlapping set of neurons (70%) contributes to line attractor dynamics during watching or engaging in fighting (Extended Data 2a-g).

While these observed attractor dynamics could be intrinsic, they might also arise from unmeasured ramping sensory input or dynamics inherited from another brain region. Although behavioral perturbations in prior studies have hinted at the intrinsic nature of VMHvl line attractor dynamics^9^, a rigorous test requires manifold-level perturbations^30 31^ targeted to cells identified as contributing to the attractor. Direct on-manifold perturbation has previously been performed only in the *Drosophila* ring attractor system^7,18^; moreover off-manifold perturbations were not performed. In mammals, although a point attractor has been perturbed using optogenetic manipulation^14,15,24^, direct single-cell perturbations of neurons contributing to a continuous attractor *in vivo* has not been reported.

To address this, we employed 2-photon (2P) imaging in head-fixed mice expressing both jGCaMP7s^32^ and ChRmine^33^ (a red-shifted opsin) during aggression observation (Figure 1a-c). rSLDS analysis identified an integration dimension with slow dynamics (x_1_) aligned to a line attractor and an orthogonal dimension with faster dynamics (x_2_) (Figure 1d-g). In neural state space, activity entered the attractor following removal of the demonstrator mice, decaying according to the system’s intrinsic leak rate (Figure 1h). We used the mapping between neural activity and the underlying state space to directly identify and image neurons contributing to each dimension. Neurons contributing to the integration dimension displayed more persistence than those aligned with the faster dimension (Figure 1i-m). Thus, a line attractor can be recapitulated in head-fixed mice observing aggression, opening the way to 2-photon-based perturbation experiments.

**Figure 1.**
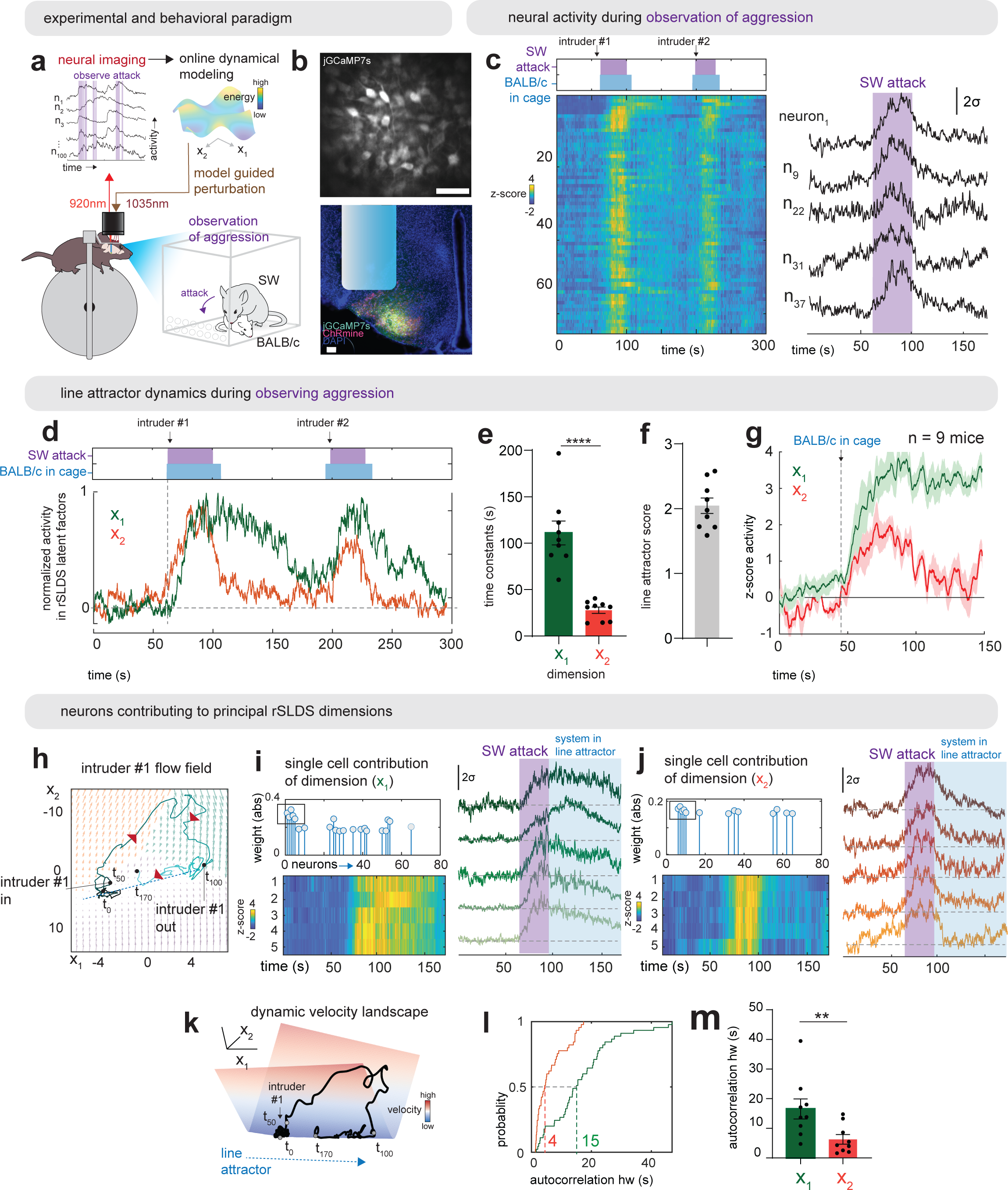
Attractor dynamics in head-fixed mice observing aggression. a. Experimental paradigm for 2-photon head-fixed mice observing aggression. 920nm 2-photon laser was used to monitor activity of VMHvl^Esr1^ neurons in head-fixed mice observing aggression. A dynamical model fit to neural data guided holographic activation of specific neurons expressing ChRmine using a 1035nm 2-photon laser. b. Example field of view in 2-photon setup through a GRIN lens (top). Fluorescence image of a coronal slice showing expression of jGCaMP7s and ChRmine (bottom). Scale bars – 100µm. c. Neural and behavioral raster from example mouse observing aggression in the 2-photon setup. Arrows indicate insertion of submissive BALB/c intruder to the observation chamber for interaction with an aggressive Swiss Webster mouse (SW). Right: Example neurons from left. d. Neural activity projected onto rSLDS dimensions obtained from models fit to 2-photon imaging data in one example mouse. e. rSLDS time constants across mice (n = 9 mice, ****p<0.0001). f. Line attractor score (see methods) across mice (n = 9 mice). g. Behavior triggered average of x_1_ and x_2_ dimensions, aligned to introduction of BALB/c into resident’s cage (n = 9 mice). Dark line – mean activity, shaded surrounding – sem. h. Flow fields from 2P imaging data during observation of aggression from one example mouse. Red arrows indicate the direction flow of time. i. Top: Identification of neurons contributing to x_1_ dimension from rSLDS model. Weight of each neurons shown as absolute value. Bottom: Activity heatmap of five neurons contributing most strongly to x_1_ dimension. Right: Neural traces of the same neurons and an indication of when the systems enters the line attractor j. Same as i but for x_2_ dimension. k. Dynamic velocity landscape from 2P imaging data during observation of aggression from one example mouse. Blue color reflects stable area in the landscape, red – unstable. Black line is the trajectory of the neuronal activity. l. Cumulative distributions of autocorrelation half width of neurons contributing to x_1_ (green) and x_2_ (red) dimensions (n = 9 mice, 45 neurons each for x_1_ and x_2_ distributions). m. Mean autocorrelation half width across mice for neurons contributing to x_1_ and x_2_ dimensions (n = 9 mice, **p<0.01).

### Holographic activation reveals the intrinsic nature of VMHvl line attractor dynamics

Next, to determine whether VMHvl^Esr1^ line attractor dynamics are intrinsic, we performed holographic activation of a subset of neurons contributing to the integration dimension (x_1_). These neurons were identified in real-time using rSLDS fitting of data recorded during observation of aggression, followed by reactivation of those neurons (in a manual closed-loop) after removing the demonstrator mice. Approximately 25% of integration-dimension neurons in the observation field were reactivated during each trial (5 cells/trial). Repeated pulses of optogenetic stimulation (2 sec, 20 Hz, 5 mW/mm^2^) were delivered with a 20s inter-stimulus interval (ISI) (Figure 2a, e).

**Figure 2.**
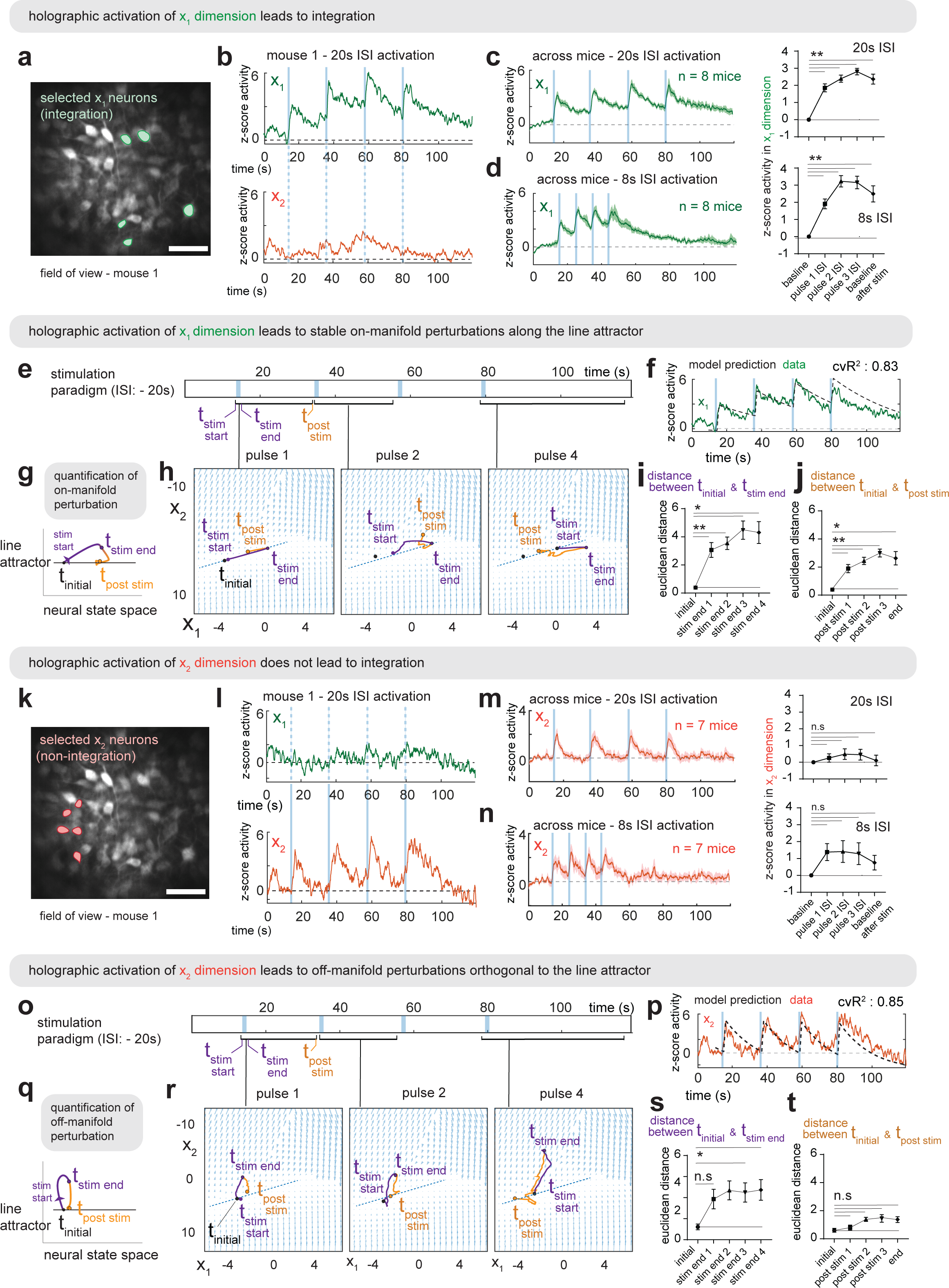
Holographic perturbations reveal line attractor dynamics in the VMHvl. a. Field of view of five x_1_ neurons selected for 2-photon activation in example mouse 1. b. Neural activity projected onto x_1_ dimension after grouped optogenetic activation of five x_1_ neurons in example mouse 1 (top). Neural activity projected onto x_2_ dimension from activation of the same x_1_ neurons (bottom). Blue vertical lines indicate the time of holographic activation of x_1_ neurons. Dashed blue lines indicate the time of holographic activation on non-activated x_2_ neurons. c. Left: Average activity projected onto x_1_ dimension from activation of x_1_ neurons across mice using 20s inter stimulus interval (n = 8 mice). Right: Quantification of average z-scored activity of projected x_1_ dimension during baseline or various inter stimulus intervals (n = 8 mice, **p<0.01). d. Same as “c” but for 8s inter stimulus interval. e. Stimulation paradigm for grouped activation of x_1_ neurons using 20s ISI. f. Data and model prediction of applying stimulation paradigm in 2e to rSLDS model trained on observation of aggression. g. Cartoon showing quantification of perturbation along line attractor in neural state space. The Euclidian distance between time points t_initial_ (baseline before stimulation) and t_stim end_ (end of stimulation) as well as between t_initial_ and t_post stim_ (end of ISI of stimulation) are calculated. h. Flow fields from example mouse 1, showing perturbations along line attractor upon activation of x_1_ neurons. i. Euclidian distance between time points t_initial_ and t_stim end_ across mice (*p<0.05, **p<0.01). j. Same as 2i but for time points t_initial_ and t_post stim_ across mice (*p<0.05, **p<0.01). k. Field of view of five x_2_ neurons selected for activation in example mouse 1. l. Neural activity projected onto x_1_ dimension after grouped optogenetic activation of x_2_ neurons in example mouse 1 (top). Neural activity projected onto x_2_ dimension from activation of same x_1_ neurons (bottom). m. Left: Average of activation of x_2_ neurons across mice using 20s inter stimulus interval (n = 8 mice). Right: Quantification of average z-scored activity during baseline or inter stimulus intervals (n.s, n = 8 mice). n. Left: Average activity projected onto x_2_ dimension from activation of x_2_ neurons across mice using 8s inter stimulus interval (n = 7 mice). Right: Quantification of average z-scored activity of projected x_2_ dimension during baseline or inter stimulus intervals (n.s, n = 7 mice). o. Stimulation paradigm for grouped activation of x_2_ neurons using 20s ISI. p. Data and model prediction of applying stimulation paradigm in 2o to rSLDS model trained on observing aggression. q. Same as 2g but for an example perturbation orthogonal to a line attractor. r. Flow fields from example mouse 1, showing perturbations orthogonal to line attractor upon activation of x_2_ neurons. s. Same as 2i but for x_2_ activation. t. Same as 2j but for x_2_ activation.

Under these conditions, activity along the x_1_ (but not the x_2_) dimension is expected to integrate, based on the time constants of these dimensions extracted from the fit rSLDS model (Figure 1e). Consistent with this expectation, optogenetic re-activation of x_1_ neurons yielded robust integration along the x_1_ dimension, as evidenced by progressively increasing activity during the ISI following each consecutive pulse (Figure 2b-c; n=8 mice). Activated x_1_ neurons exhibited activity levels comparable to their response during observation of aggression (Extended Data 3a-c). Similar results were obtained using an 8s ISI (Figure 2d). Providing the same input to the fit rSLDS model also resulted in integration along the x_1_ dimension similar to that observed in the data (Figure 2f). Importantly, x_1_ stimulation did not evoke discernable activity in the x_2_ dimension, suggesting a lack of functional interaction between these dimensions at a population level (Extended Data 3d-f and see below).

To visualize the effect of re-activation of x_1_ neurons in neural state space, we projected the data into a 2D flow-field based on the first two PCs of the reduced rSLDS space. Activation pulses transiently moved the neural population vector “up” the line attractor, followed by relaxation back down the attractor to a point that was higher than the initial position of the system (Figure 2h, pulse 1, 2 & 4). To quantify this effect, we calculated the Euclidean distance in state space between the initial time point during the baseline period (denoted t_initial_), to the time point at the end of stimulation or at the end of the ISI following each pulse (denoted t_stim end_ and t_post stim_ respectively) (Figure 2g). This revealed that the x_1_ perturbations resulted in progressive, stable on-manifold movement along the attractor with each consecutive stimulation, as measured by the increase in both metrics (Figure 2h-j).

Importantly, as predicted by rSLDS, activation of x_2_ neurons did not lead to integration (Figure 2k, l). Instead, following each pulse we observed stimulus-locked transient activity in the x_2_ dimension followed by a decay back to baseline during the ISI period, across stimulation paradigms (Figure 2k-n), with little to no effect on x_1_ neurons (Extended Data 3g-i). In 2D neural state space, we observed that x_2_ neuron activation caused transient off-manifold movements of the population activity vector orthogonal to the attractor axis during each pulse (Figure 2q-s). Following each stimulus, the neural trajectory relaxed back into the attractor, at the initial location it occupied before the stimulus. The small Euclidean distance between t_initial_ and t_post stim_ underscored the attractor’s stability (Figure 2t). Activation of randomly selected neurons not weighted by either dimension did not produce activity along either the x_1_ or x_2_ dimension, emphasizing the specificity of our on- and off-manifold holographic activation (Extended Data 3j-n). Taken together, these findings provide evidence for the intrinsic nature of the VMHvl^Esr1^ line attractor.

### Line attractor-contributing neurons form selective functional ensembles

Line attractors have traditionally been hypothesized to emerge from recurrent interactions within a network (although single neurons can also have the potential to integrate using neuromodulator regulated ion channels)^34^. To determine whether network-level interactions contribute to the implementation of the line attractor, we performed single-cell activation of either individual x_1_ or x_2_ neurons combined with imaging of unperturbed neurons, to assess functional connectivity within the circuit (Figure 3a). These experiments revealed selective functional coupling between x_1_ neurons, as evidenced by an increase in activity during the ISI period in unperturbed x_1_ neurons following each pulse of activation (Figure 3b, d). However, we observed little activity in unperturbed x_2_ neurons upon activation of single x_1_ neurons (Figure 3c, e), indicating that functional x_1_ connectivity is selective.

**Figure 3.**
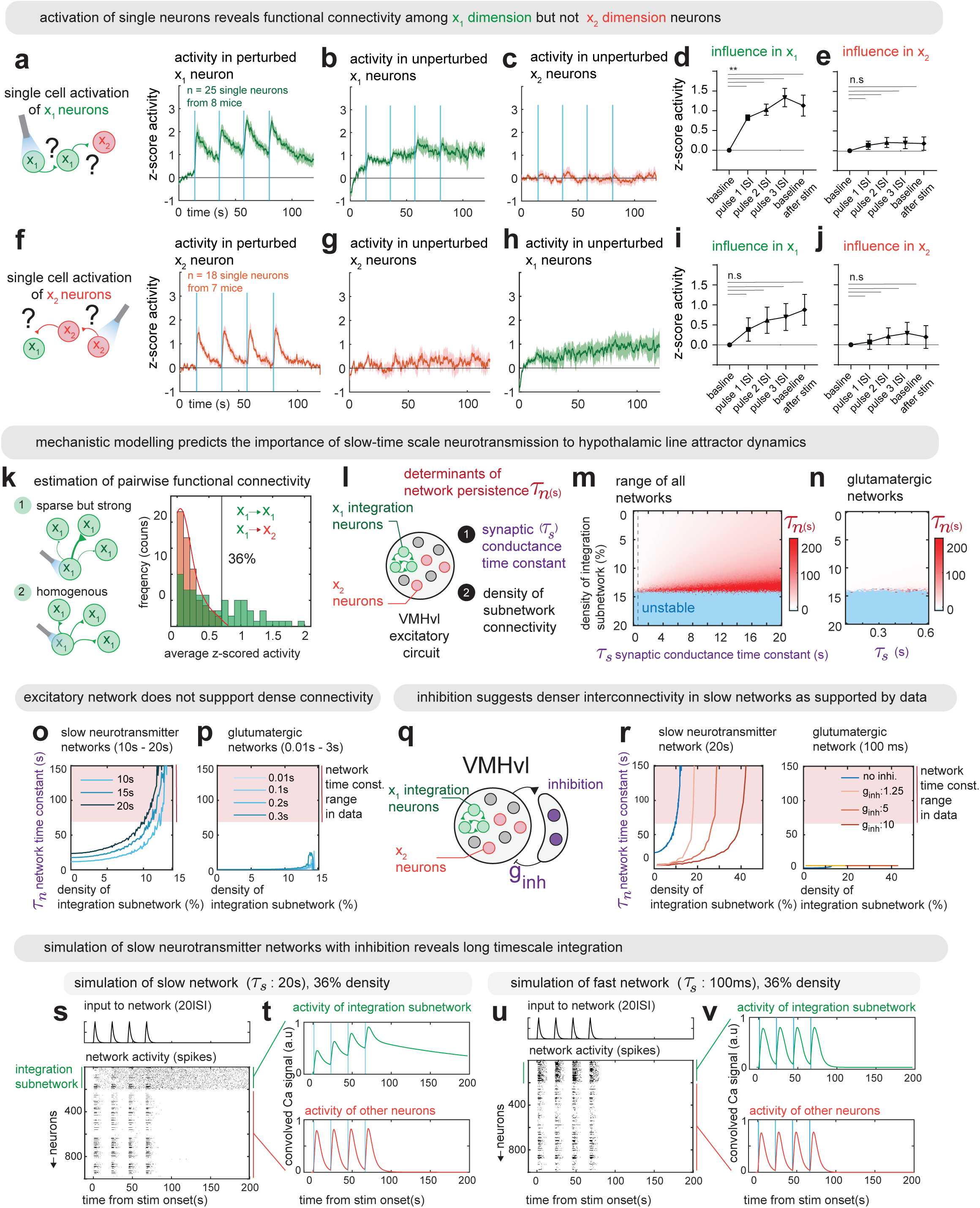
Neural implementation of a line attractor guided by functional connectivity. a. Left: Paradigm for examining activity in unperturbed x_1_ and x_2_ neurons upon activation of single x_1_ neurons. Right: Average z-score activity of perturbed x_1_ neurons (25 single neurons from n = 8 mice). b. Average z-score activity of unperturbed x_1_ neurons upon perturbation of single x_1_ neurons (n = 8 mice). c. Average z-score activity of unperturbed x_2_ neurons upon perturbation of single x_1_ neurons (n = 8 mice). d. Quantification of activity in unperturbed x_1_ neurons upon perturbation of single x_1_ neurons (**p<0.01, n = 8 mice). e. Quantification of activity in unperturbed x_1_ neurons upon perturbation of single x_2_ neurons (n.s, n = 8 mice). f. Left: Paradigm for examining activity in unperturbed x_1_ and x_2_ neurons upon activation of single x_2_ neurons. Right: Average z-score activity of perturbed x_2_ neurons (18 single neurons from n = 7 mice). g. Average z-score activity of unperturbed x_2_ neurons upon perturbation of single x_2_ neurons (n = 7 mice). h. Average z-score activity of unperturbed x_1_ neurons upon perturbation of single x_2_ neurons (n = 7 mice). i. Quantification of activity in unperturbed x_1_ neurons upon perturbation of single x_2_ neurons (n.s, n = 7 mice). j. Quantification of activity in unperturbed x_2_ neurons upon perturbation of single x_2_ neurons (n.s, n = 7 mice). k. Left: Cartoon illustrating either strong but sparse connectivity among x1 neurons (1), or dense and interconnectivity within subnetwork (2). Right: Empirical distribution of pairwise functional connectivity between x_1_ neurons (green) and from x_1_ to x_2_ neurons (red) (n = 99 pairs, n = 7 mice). l. Cartoon illustrating different elements of an excitatory network that can determine network level persistent activity including synaptic conductance time constant (t_s_) and density of subnetwork connectivity (σ). m. Model simulation result showing network time constant (t_n_) by varying σ in range 0-20% density values and t_s_ in range 0-20s. The blue portions of the image refer to configurations that result in unstable networks with runaway excitation. n. Zoomed in version of 3m but for glutamatergic networks with synaptic conductance time constant (t_s_) in range 0.01-0.5s. o. Plot of network time constant (t_n_) against density of integration subnetwork for slow neurotransmitter networks (t_s_:10,15,20s). The network time constant t_n_ varies monotonically with density for large values of t_s_. p. Same as 3o but for glutamatergic networks (t_s_:0.01,0.1,0.2,0.3s). q. Cartoon showing modified VMHvl circuit with fast feedback inhibition incorporated. r. Left: Plot of network time constant (t_n_) against density of integration subnetwork for a slow neurotransmitter network with t_s_ = 20s, for different values of strength of inhibition (inhibitory gain, g_inh_: 1.25,5,10). Right: Same as left but for a glutamatergic network with t_s_= 0.1s. s. Model simulation of a slow neurotransmitter network with fast feedback inhibition (t_s_:20s, 36% density of subnetwork connectivity). Top: Input (20s ISI) provided to model, Bottom: Spiking activity in network. The first 200 neurons (20%) comprise the interconnected integration subnetwork. t. Ca^2+^ activity convolved from firing rate (see Methods) of integration subnetwork (top) and remaining neurons (bottom). u. Same as 3s but for a fast transmitter network (t_s_:0.1s, 36% density of subnetwork connectivity). v. Same as 3t but for a fast transmitter network (t_s_:0.1s, 36% density of subnetwork connectivity).

In contrast, activation of single x_2_ neurons revealed a lack of functional coupling between x_2_ neurons (Figure 3g). While there was some increase in activity in unperturbed x_1_ neurons upon activation of single x_2_ neurons (Figure 3h), that increase was not significant, suggesting that x_2_ neurons might not be coupled with other x_1_ or x_2_ neurons (Figure 3i, j).

The functional connectivity we observed could arise either from a population of sparsely but strongly inter-connected neurons, or from a population with denser connections of intermediate strength^35^ (Figure 3k, left). To assess this, we calculated the distribution of pairwise influences, defined as the average evoked activity in each unperturbed integration neuron post stimulus. To estimate an upper bound on the amount of influence within the network, we considered the percentage of integration pairs that had influence scores higher than the maximum influence of x_1_ onto x_2_ neurons (Figure 3k, right). This analysis revealed a densely connected integration subnetwork with a connection density of about 36% (Figure 3k, right). These data suggest that VMHvl^Esr1^ neurons that contribute to the line attractor form functional ensembles, confirming theory-based predictions^34^.

We next used computational approaches to investigate the nature of the observed functional connectivity within x_1_ ensembles. Such connectivity could reflect different types of synapses: they could be fast and glutamatergic, as typically assumed for most attractor networks^34^; or they could be slow neuromodulator-based connections that use GPCR-mediated second messenger pathways to sustain long time-scale changes in synaptic conductance. To investigate systematically the density and synaptic kinetics of networks capable of generating line attractors with the observed integration-dimension (x_1_) network time constant, we turned to mechanistic modelling using an excitatory integrate and fire network^36^ (Figure 3l). Because VMHvl is exclusively glutamatergic^37^, we used excitatory networks and analytically calculated the network time constant using an eigen-decomposition of the connectivity matrix^34^ (Extended Data 4a). By varying the synaptic conductance time constant (τ_*s*_) and the density of integration subnetwork connectivity (σ), we found that only artificial networks based on relatively sparse connectivity (∼8-12%) and slow synaptic time constants (20s) could yield network time constants (τ_*n*_) in the experimentally observed range (∼50-200s; Figure 3m, o; red shading). In contrast, networks with fast glutamatergic connectivity failed to do so over the same range of connection densities (Figure 3n, p).

In these purely excitatory network models, the density of connections that yielded network time constants in the observed range was much lower than the experimentally measured value (36%). To match more accurately the empirically observed connection density, we incorporated excitation-recruited fast-feedback inhibition into our integrate- and-fire network^36^; VMHvl is known to receive dense GABAergic innervation from surrounding areas^38^. The addition of global strong feedback inhibition allowed networks to match the observed connection density (36%), but importantly, maintained the slow nature of the functional connectivity (20s; Figure 3q, r). Indeed, networks simulated with a long τ_*s*_ (20s) and dense σ (36%) could integrate digital optogenetic stimulation in a manner similar to that observed experimentally (Figure 3s-t; cf. 3a). In contrast, purely glutamatergic networks (τ_*s*_=100 msec) were unable to integrate at the observed timescales (Figure 3u-v). Together, these results suggest an implementation of the VMHvl^Esr1^ line attractor that combines slow neurotransmission and dense subnetwork interconnectivity within an attractor creating ensemble.

### Strength of functional connectivity correlates with attractor stability across mice

The stability of the line attractor in VMHvl during aggression has previously been positively correlated with aggressiveness across individual mice^9^. We therefore investigated whether individual mouse differences in the stability of the line attractor might be correlated with the strength of functional connectivity within the x_1_ ensemble (Figure 4a). To do so, we compared the x_1_ time constants from rSLDS models fit to imaging data recorded during attack observation from each mouse (a measure of attractor stability) to different quantitative metrics of functional connectivity strength (such as average z-scored activity or area under the curve), measured by optogenetic stimulation of single x_1_ or x_2_ neurons and imaging of unperturbed cells in the same animals following removal of the demonstrator intruder mice (Figure 4b, c).

**Figure 4.**
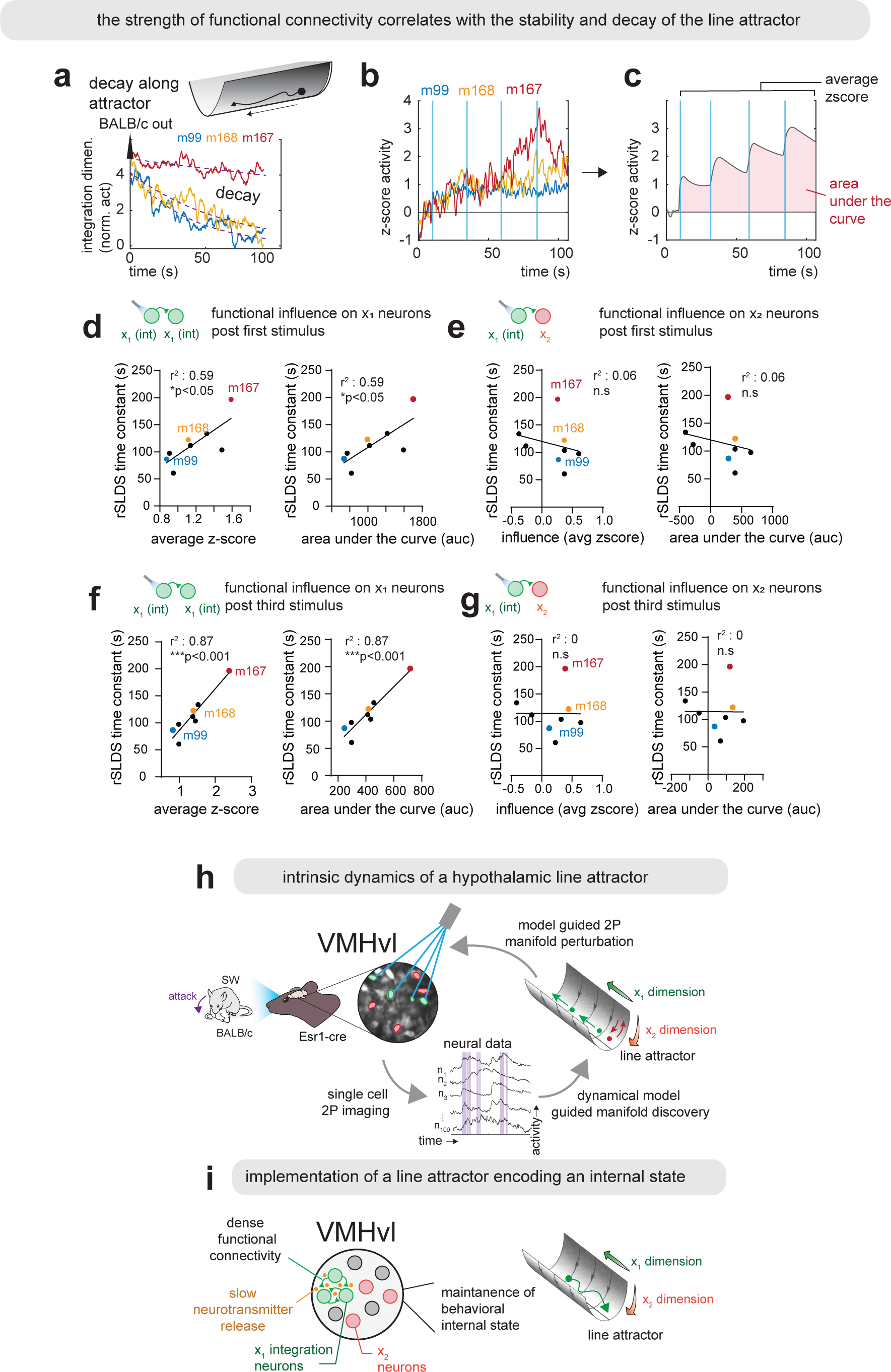
Strength of functional connectivity infers line attractor’s decay rate. a. Example neural activity projected onto x_1_ (integration) dimension of three mice that display varying rates of decay following removal of a Balb/c intruder from cage while observing aggression (i.e movement down the line attractor). b. Z-score activity on unperturbed x_1_ neurons upon activation of single x_1_ neurons in the same mice from 4a. c. Illustration of different quantification approaches to the change in activity of unperturbed x_1_ neurons from 4b as either mean z-score activity following first stimulus or area under the curve (auc). d. Correlation between rSLDS time constant obtained from observation of aggression and average z-score across unperturbed x_1_ neurons across 8 mice measured using either average z-score (left) or AUC (right) post first stimulus (r^2^: 0.59, *p<0.05). e. Correlation between rSLDS time constant obtained from observation of aggression and average z-score across unperturbed x_2_ neurons across 8 mice measured using either average z-score (left) or AUC (right) post first stimulus (r^2^: 0.06, n.s). f. Same as 4d but quantified post third stimulus. (r^2^: 0.87, ***p<0.001) g. Same as 4e but quantified post third stimulus. (r^2^: 0.0, n.s) h. Cartoon depicting summary of results illustrating intrinsic dynamics of hypothalamic line attractor. i. Cartoon depicting implementation of a hypothalamic line attractor encoding a behavioral internal state.

Remarkably, there was a strong correlation across mice between the time constant of the line attractor measured during the observation of aggression, and the strength of functional connectivity among integration-dimension (x_1_) neurons measured by post-observation optogenetic stimulation (Figure 4d). The strength of this correlation was higher after the third (r^2^=0.87) than the first (r^2^=0.59) stimulus, suggesting that individual differences in integration dynamics become more apparent once the system has already integrated several inputs (Figure 4f). Importantly, this correlation was specific to functional connectivity within the integration subnetwork and did not hold when rSLDS time constants were compared with the influence strength of stimulated x_1_ neurons on x_2_ cells (Figure 4e, g). Thus, individual differences among mice in the stability of the line attractor during the observation of aggression are correlated with differences in the functional connection strength among attractor-contributing neurons, as measured post-observation by optogenetic stimulation and imaging of the same cells in the same animals.

## Discussion

Using model-guided closed-loop all-optical experiments, we have directly demonstrated line attractor dynamics in a mammalian system (Figure 4h, i). Attractors are intrinsic properties of neural networks that emerge from network interactions. To distinguish such intrinsic properties from the inheritance of attractor-like dynamics from upstream inputs, specific neural perturbations are essential. Perturbations using bulk optogenetics and electrophysiological recording have been pivotal in demonstrating point attractor dynamics encoding short term memory in the mammalian brain^14^. While such experiments can illuminate important features of network stability, our experiments used both on- and off-manifold 2-photon optogenetic perturbations at single-cell resolution, in combination with calcium imaging, to definitively test the intrinsic nature of a continuous line attractor that was initially discovered by rSLDS modeling. While on-manifold perturbations were used previously to experimentally move neural activity along a ring attractor encoding head-direction in *Drosophila*, off-manifold perturbations demonstrating the key property of “attractiveness” were not performed in that case^18,19^. To our knowledge, therefore the present results constitute the first *in vivo* on- and off-manifold perturbation experiments demonstrating the intrinsic properties of a continuous attractor in any system.

Our data and modeling also provided insight into the implementation of the line attractor. We found evidence of dense, specific subnetwork connectivity coupled with slow neurotransmission. Although our models confirm the importance of rapid feedback inhibition, as indicated in invertebrate ring attractor studies^18^, they diverge markedly from conventional continuous attractor models^4,34^ by highlighting the role of slow neurotransmission over rapid excitation. Numerous theoretical studies have posited that continuous attractors relying on fast recurrent connectivity are not robust due to the necessity for precise tuning of synaptic weights to sustain stable attractor dynamics^34,39^. The slow neurotransmission predicted by our model could be implemented by GPCR-mediated signaling triggered by biogenic amines or neuropeptides^40^. Consistent with this prediction, we have recently found that VMHvl line attractor dynamics are dependent on signaling through oxytocin and/or vasopressin neuropeptide receptors expressed in Esr1^+^ neurons^41^. These findings suggest an evolutionary mechanism that favors the robustness offered by slow neurotransmission in the establishment of a line attractor encoding a persistent internal motive state. Whether line attractors in other systems that mediate cognitive functions on shorter time-scales^11,42^ indeed rely primarily on fast glutamatergic recurrent connectivity remains to be determined.

Lastly, our observations indicate a pronounced correlation between individual differences in the functional strength of integration subnetwork connectivity, as revealed by optogenetic stimulation and imaging, and differences in the measured stability of the line attractor evoked by a naturalistic stimulus. This suggests that attributes of the attractor, such as its connectivity density or strength, may be modifiable (either by genetics and/or experience^43^), and may underlie individual differences in aggressiveness^9^. Deciphering the underlying mechanisms that grant this attractor its apparent flexibility represents a promising avenue for future research.

---

**Extended Data 1.**
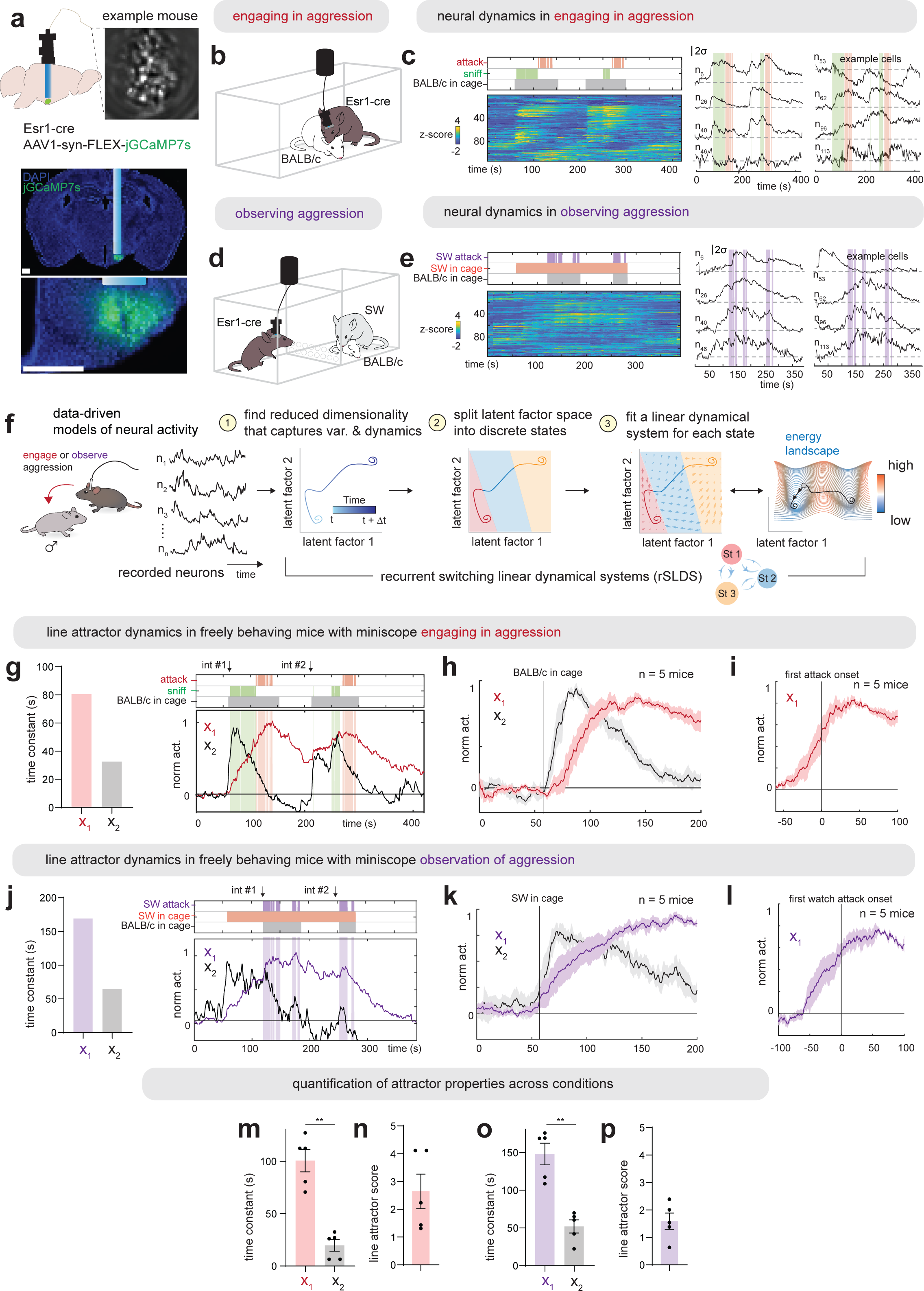
Shared line attractor dynamics in engaging and observing aggression. a. Implantation of miniscope, field of view (top) and fluorescence image showing histology (bottom) with jGCaMP7s expression in VMHvl. b. Experimental paradigm to record VMHvl^Esr1^ activity in mice engaging in aggression. c. Left: neural & behavioral raster of example mouse 1 when engaging in aggression. Right: example neurons. d. Experimental paradigm to record VMHvl^Esr1^ activity in same mice in Ex. Data 1c during observation of aggression. e. Left: neural & behavioral raster of example mouse 1 during observation of aggression. Right: example neurons. f. Overview of rSLDS analysis. g. Left: rSLDS time constants in example mouse 1. Right: Neural activity projected onto two dimensions (x_1_ & x_2_) of dynamical system. h. Behavior triggered average of x_1_ and x_2_ dimensions, aligned to introduction of male intruder (n = 5 mice) i. Behavior triggered average of x_1_ dimensions, aligned to first attack onset (n = 5 mice). j. Left: rSLDS time constants in example mouse 1 during observation of aggression. Right: Neural activity projected onto two dimensions (x_1_ & x_2_) of dynamical system. k. Behavior triggered average of x_1_ and x_2_ dimensions from observation of aggression, aligned to introduction of Balb-c into resident’s cage (n = 5 mice). l. Behavior triggered average of x_1_ dimensions from observation of aggression, aligned to first bout of watching attack (n = 5 mice). m. rSLDS time constants across mice engaging in aggression (n = 5 mice). n. Line attractor score across mice engaging in aggression (n = 5 mice). o. rSLDS time constants across mice during observation of aggression (n = 5 mice). p. Line attractor score across mice during observation of aggression (n = 5 mice).

**Extended Data 2.**
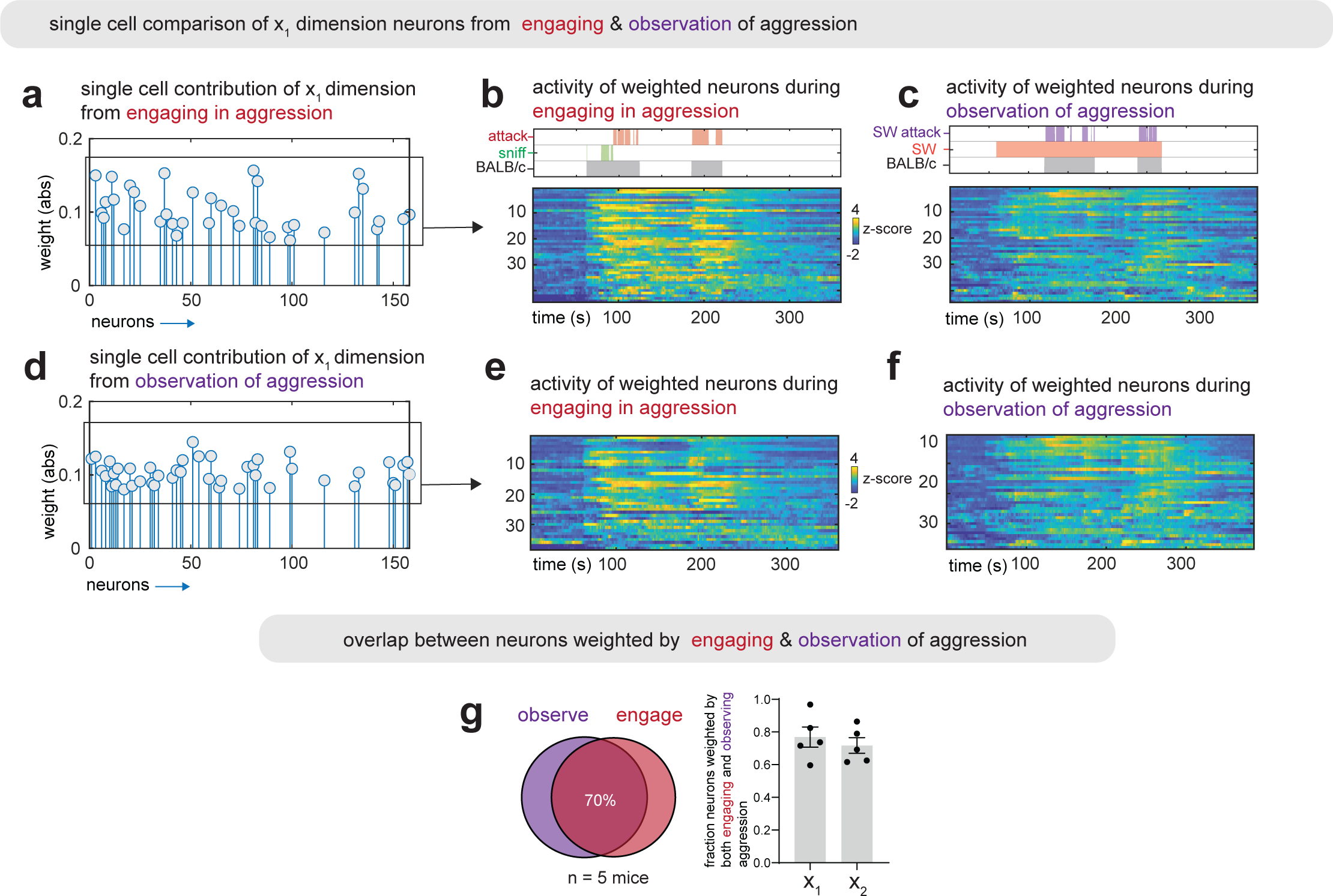
Single cell comparison of integration neurons across conditions. a. Single cell contribution of x_1_ dimension (rSLDS weights) from engaging in aggression in example mouse. b. Z-score activity of weighted neurons from Ex. Data 2a during engaging in aggression from same mouse. c. Z-score activity of weighted neurons from Ex. Data 2a during observation of aggression from same mouse. d. Single cell contribution of x_1_ dimension (rSLDS weights) from observation of aggression in example mouse. e. Z-score activity of weighted neurons from Ex. Data 2d during engaging in aggression from same mouse. f. Z-score activity of weighted neurons from Ex. Data 2d during observation of aggression from same mouse. g. Overlap in neurons contributing to line attractor (x_1_) & x_2_ dimension from rSLDS performing independently in engaging versus observing aggression. Left: Example mouse, Right: Average across 5 mice.

**Extended Data 3.**
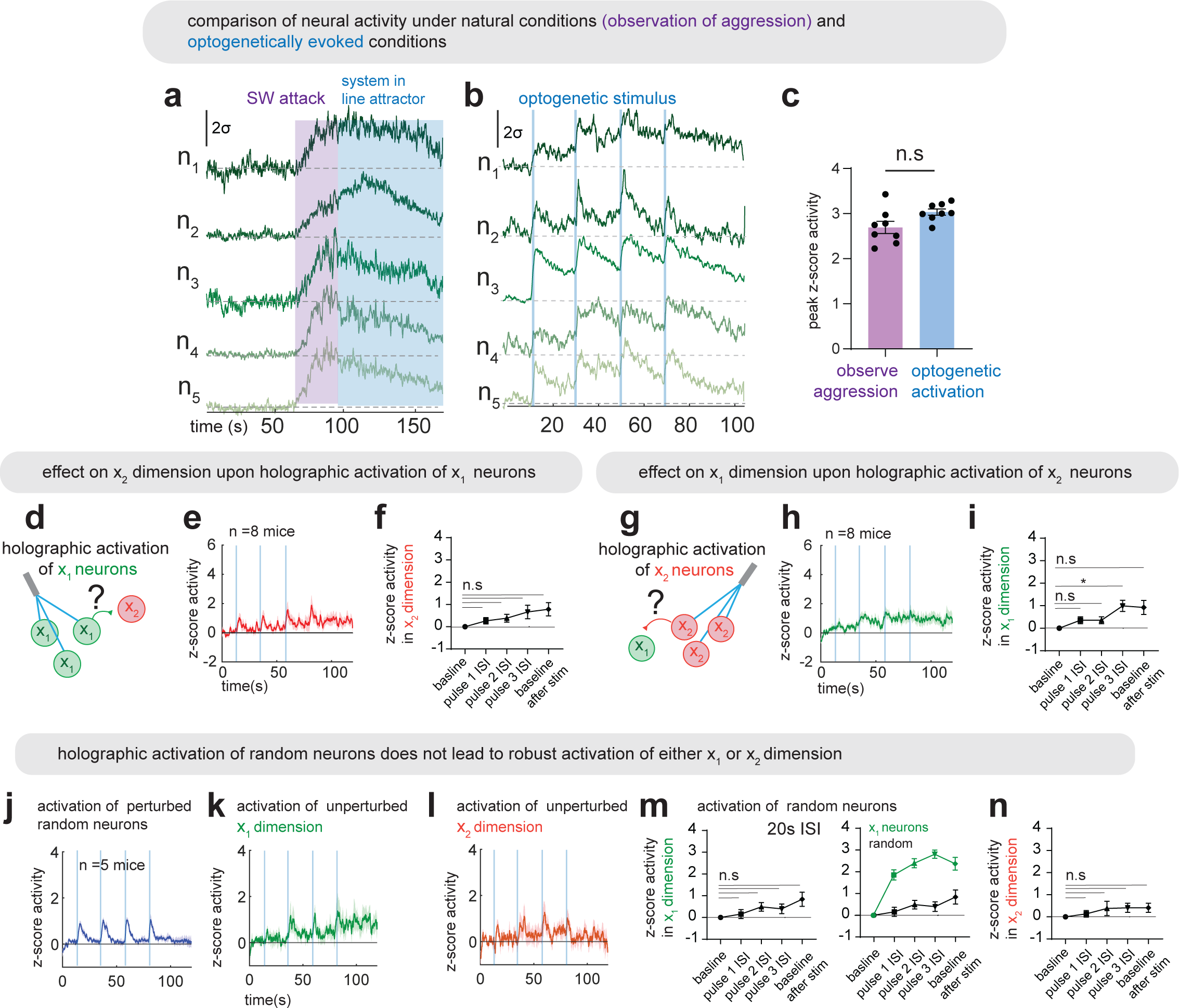
Interaction between dimensions and activation of random neurons. a. Neural activity of five x_1_ neurons selected for grouped optogenetic perturbation during observation of aggression. b. Neural activity of same five x_1_ neurons in Ex. Data 3a during grouped optogenetic activation. c. Comparison of peak z-score of x_1_ neurons selected for grouped optogenetic activation during observation of aggression and during optogenetic activation (n = 9 mice). d. Paradigm for examining activity in x_2_ dimension upon grouped holographic activation of x_1_ neurons. e. Average z-score activity of neural activity projected onto x_2_ dimension across mice (n = 8 mice). f. Quantification of activity in unperturbed x_2_ dimension upon grouped holographic activation of x_1_ neurons (n.s, n = 8 mice). g. Paradigm for examining activity in x_1_ dimension upon grouped holographic activation of x_2_ neurons. h. Average z-score activity of neural activity projected onto x_1_ dimension across mice (n = 8 mice). i. Quantification of activity in unperturbed x_1_ dimension upon grouped holographic activation of x_2_ neurons (*p<0.05, n = 8 mice). j. Effect of grouped holographic activation of randomly selected neurons on activated neurons. k. Average z-score activity of unperturbed x_1_ dimension upon activation of random neurons (n = 5 mice). l. Average z-score activity of unperturbed x_2_ dimension upon activation of random neurons (n = 5 mice). m. Left: Quantification of activity in unperturbed x_1_ dimension upon grouped holographic activation of random neurons (n.s, n = 5 mice). Right: Comparison of grouped activation of x_1_ neurons (green, reproduced from Fig. 2c, right) and grouped activation of random neurons on activity of x_1_ dimension (black, reproduced from Ex. Data 3m, left). n. Quantification of activity in unperturbed x_2_ dimension upon grouped holographic activation of random neurons (n.s, n = 5 mice).

**Extended Data 4.**
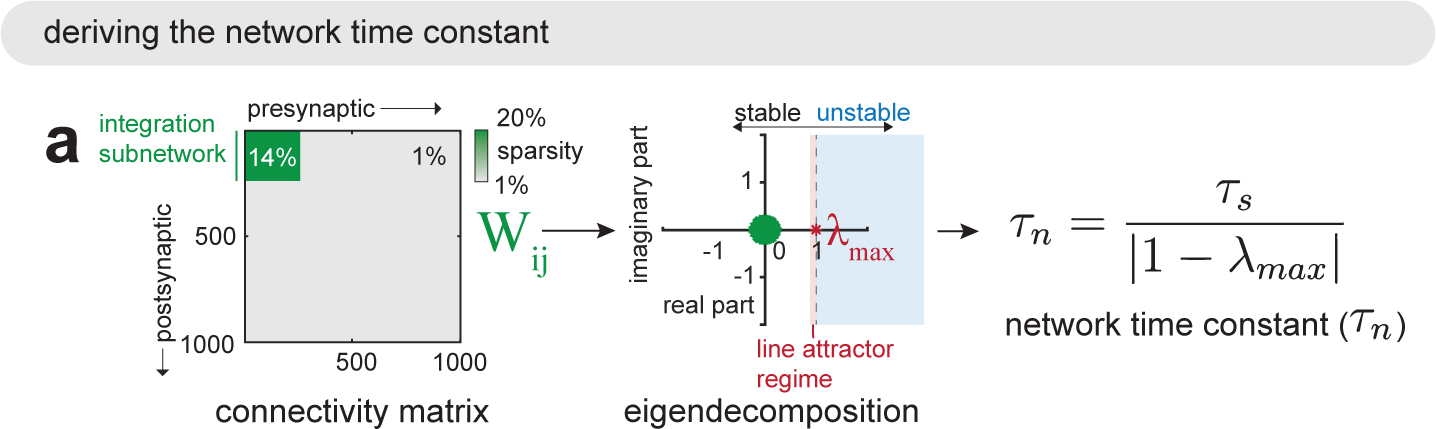
Deriving network time constant for model simulations. a. Analytical derivation of network time constant from connectivity matrix of purely excitatory recurrent neural network.

## Methods

### Mice

All experimental procedures involving the use of live mice, or their tissues were carried out in accordance with NIH guidelines and were approved by the Institute Animal Care and Use Committee and the Institute Biosafety Committee at the California Institute of Technology (Caltech). All C57BL/6N (Bl/6N) mice used in this study, including wild-type and transgenic mice, were bred at Caltech. Swiss Webster (SW) male Residents and BALB/c male intruder mice were bred at Caltech. Experimental Bl/6N mice and resident SW mice were used at the age of 8–20 weeks. Intruder BALB/c mice were used at the age of 6–12 weeks and were maintained with three to five cage mates to reduce their aggression. Esr1Cre/+ knock-in mice (Jackson Laboratory, stock no. 017911) were back-crossed into the Bl/6N background (>N10) and bred at Caltech. Heterozygous Esr1Cre/+ mice were used for cell-specific targeting experiments and were genotyped by PCR analysis using genomic DNA from ear tissue. All mice were housed in ventilated micro-isolator cages in a temperature-controlled environment (median temperature 23 °C, humidity 60%), under a reversed 11/13-h dark/light cycle, with ad libitum access to food and water. Mouse cages were changed weekly.

### Viruses

The following adeno-associated viruses (AAVs), along with the supplier, injection titers (in viral genome copies ml–1 (vg ml–1) and injection volumes (in nanoliters), were used in this study: AAV1-syn-FLEX-jGCaMP7s-WPRE (Addgene, no. 104492, roughly 2 × 10^12^ vg ml–1, 200 nl per injection), AAVdj-Ef1a-DIO-ChRmine-mScarlet-Kv2.1-WPRE (Janelia Vector Core, around 2 × 10^12^ vg ml–1, 200 nl per injection).

### Histology

Following completion of 2-photon\miniscope experiments, histological verification of virus expression and implant placement were performed on all mice. Mice lacking virus expression or correct implant placement were excluded from the analysis. Mice were perfused transcardially with 0.9% saline at room temperature, followed by 4% paraformaldehyde (PFA) in 1× PBS. Brains were extracted and post-fixed in 4% PFA overnight at 4 °C, followed by 24 h in 30% sucrose/PBS at 4 °C. Brains were embedded in OCT mounting medium, frozen on dry ice and stored at −80 °C for subsequent sectioning. Brains were sectioned in 80-μm thickness on a cryostat (Leica Biosystems). Sections were washed with 1× PBS and mounted on Superfrost slides, then incubated for 30 min at room temperature in DAPI/PBS (0.5 μg/ml) for counterstaining, washed again and coverslipped. Sections were imaged with epifluorescent microscope (Olympus VS120).

### Stereotaxic Surgeries

Surgeries were performed on sexually experienced adult male Esr1Cre/+mice aged 6– 12 weeks. Virus injection and implantation were performed as described previously^22,44^. Briefly, animals were anaesthetized with isoflurane (5%for induction and 1.5% for maintenance) and placed on a stereotaxic frame (David Kopf Instruments). Virus was injected into the target area using a pulled-glass capillary (World Precision Instruments) and a pressure injector (Micro4 controller, World Precision Instruments), at a flow rate of 50 nl min-1. The glass capillary was left in place for 5 min following injection before withdrawal. Stereotaxic injection coordinates were based on the Paxinos and Franklin atlas^45^. Virus injection: VMHvl, AP: −1.5, ML: ±0.75, DV: −5.75. For 2-photon experiments GRIN lenses (0.6 × 7.3 mm, Inscopix) were slowly lowered into the brain and fixed to the skull with dental cement (Metabond, Parkell). Coordinates for GRIN lens implantation: VMHvl: AP: −1.5, ML: −0.75, DV: −5.55). A permanent head-bar was attached to the skull with Secure Resin cement (parkell). For micro-endoscope experiments an additional baseplate was attached to the skull (Inscopix).

### Housing conditions for behavioral experiments

All male Bl/6N mice used in this study were socially and sexually experienced. Mice aged 8–12 weeks were initially co-housed with a female Bl/6N female mouse for 1 day and were then screened for attack behaviors. Mice that showed attack towards males during a 10 min resident intruder assay were selected for surgery and subsequent behavior experiments. From this point forward, these male mice were always co-housed with a female.

### Behavior annotations

Behavior videos were manually annotated using a custom MATLAB-based behavior annotation interface^46,47^. A ‘baseline’ period of 5 min when the animal was alone in its home cage was recorded at the start of every recording session. Two behaviors during the resident intruder assays were annotated: sniff (face, body, genital-directed sniffing) towards male intruders, and attack (bite, lunge).

### Behavioral assays

An observation arena was built from a transparent acrylic (18× 12.5× 18 cm, LxWxH), and a perforated part was put in front of the mice observing aggression. Perforations were 1.27 cm diameter and spread evenly throughout the bottom third of the panel. Before initiation of the assay, the observation arena was scattered with soiled bedding from the cage of the aggressive SW demonstrator. For observation of aggression in freely behaving animals (miniscope experiments) an observer was first habituates for 15 minutes. Then, a singly housed SW male demonstrator was introduced into the observation arena, followed 1 min later with the insertion of a socially housed stimulus male (BALB/c) in the same compartment. The observation of aggressive encounters persisted for ∼1 min, then after 2 minutes a different intruder was introduced for another minute. Observation assays were conducted under white light illumination. For experiments in engaging aggression, the resident mouse was first habituated 15 minutes then a BALB/c intruder mouse was introduced twice for 1-2 minutes. For the experiments comparing neural activity of mice observing aggression and mice engaging aggression, we randomly changed the order of sessions. For mice observing aggression in the 2P setup similar the approach was similar except that the observer mouse was head-fixed and on a treadmill instead of freely behaving in his home cage.

### Micro-endoscopic imaging

On the day of imaging, mice were habituated for at least 15 min after installation of the miniscope in their home cage before the start of the behavior tests. Imaging data were acquired at 30 Hz with 2× spatial downsampling; light-emitting diode power (0.1–0.5) and gain (1–7×) were adjusted depending on the brightness of GCaMP expression as determined by the image histogram according to the user manual. A transistor–transistor logic (TTL) pulse from the Sync port of the data acquisition box (DAQ, Inscopix) was used for synchronous triggering of StreamPix7 (Norpix) for video recording.

### 2-photon imaging and holographic optogenetics

Two to three weeks after surgery mice were habituated to the experimenter’s hand by handling 15 minutes a day for three consecutive days. Once animals where habituated to the experimenter’s hand, they were manually scooped and gently placed on the treadmill. Mice were head-fixation 3 consecutive days for habituation. Head-fixation was achieved by securing the head bar into a metal clamp attached to a custom head-stage. During habituation, mice were placed underneath the objective for 15 minutes and given access to random presentations of chocolate milk. Following habituation, combined two-photon imaging and behavior sessions were conducted. jGCaMP7s imaging was acquired via an Ultima 2P Plus and the Prairie View Software (Bruker Fluorescence Microscopy, USA). Individual frames were acquired at 10Hz using a galvo-resonant scanner with a resolution of 1024px x 1024px. We used a long working distance 20x air objective designed for infrared wavelengths (Olympus, LCPLN20XIR, 0.45 numerical aperture (NA), 8.3mm working distance) combined with femtosecond-pulsed laser beam (Chameleon Discovery, Coherent). GCaMP was excited using a 920nm wavelength. For targeted photostimulation, the same microscope and acquisition system (Bruker) was used with a second laser path consisting of a 1035nm high power femtosecond pulsed laser (Monaco 1035-40-40, Coherent), spatial light modulator (512×512-pixel density) to generate multi-point stimulation montages (NeuraLight 3D, Bruker).. Neurons were selected for targeted photostimulation based on their weights from the rSLDS model. During the photostimulation sessiona 128-frame average image was generated in order to clearly highlight all neurons. Using the spiral stimulation targets (10µm diameter) were manually placed on top of GCaMP expressing neurons. Laser power was adjusted to be 1.5-5.5mW of stimulation per target. We used prairie software elicit holographic photostimulation (10hz, 2s, 10ms pulse width). Photostimulations were done between frames to avoid laser artefacts. Importantly, to reduce cross activation of the ChRmine from the 920nm laser we kept laser power for imaging to be less than 30mW.

To extract regions of interest, data from mice observing aggression was uploaded to ImageJ. Then, videos were motion corrected using the moco plugin^48^. Motion corrected videos were averaged, and additional contrast and brightness adjustments were made to clearly highlight all neurons in the field of view. Then cells were manually extracted and an rSLDS model was used to identify x_1_ and x_2_ dimension neurons. Neurons were then identified on the field of view using the prairie view software and were targeted for photo-stimulation. While rSLDS models was running (15-20 minutes, see below), control experiments were conducted.

### Micro-endoscopic data extraction

#### Preprocessing

Miniscope data were acquired using the Inscopix Data Acquisition Software as 2× downsampled .isxd files. Preprocessing and motion correction were performed using Inscopix Data Processing Software. Briefly, raw imaging data were cropped, 2× downsampled, median filtered and motion corrected. A spatial band-pass filter was then applied to remove out-of-focus background. Filtered imaging data were temporally downsampled to 10 Hz and exported as a .tiff image stack.

#### Calcium data extraction

After preprocessing, calcium traces were extracted and deconvolved using the CNMF-E^49^ large data pipeline with the following parameters: patch_dims = [4], gSig = 3, gSiz = 13, ring_radius = 17, min_corr = 0.7, min_pnr = 8. The spatial and temporal components of every extracted unit were carefully inspected manually (SNR, PNR, size, motion artefacts, decay kinetics and so on) and outliers (obvious deviations from the normal distribution) were discarded.

#### Dynamical system models of neural data

Recurrent-switching linear dynamical system (rSLDS) models^16,29^ are fit to neural data as previously described^15^. Briefly, rSLDS is a generative state-space model that decomposes non-linear time series data into a set of discrete states, each with simple linear dynamics. The model describes three sets of variables: a set of discrete states (z), a set of latent factors (x) that captures the low-dimensional nature of neural activity, and the activity of recorded neurons (y). While the model can also allow for the incorporation of external inputs based on behavior features, such external inputs were not included in our first analysis.

The model is formulated as follows: At each timepoint, there is a discrete state *z*_*t*_ ∈ {1, …, *K*} that depends recurrently on the continuous latent factors (x) as follows:

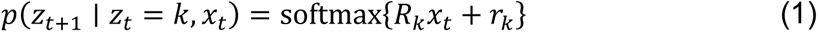

where *R*_*k*_ ∈ ℝ^*K*×*K*^ and *r*_*k*_ ∈ ℝ^*K*^ parameterizes a map from the previous discrete state and continuous state to a distribution over the next discrete states using a softmax link function. The discrete state *z*_*t*_ determines the linear dynamical system used to generate the latent factors at any time t:

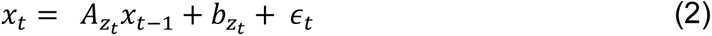

where *A*_*k*_ ∈ ℝ^*d*×*d*^ is a dynamics matrix and *b*_*k*_ ∈ ℝ^*D*^ is a bias vector, where *D* is the dimensionality of the latent space and *∈*_*t*_ ∼ *N*(0, *Q*_*zt*_) is a Gaussian-distributed noise (aka innovation) term.

Lastly, we can recover the activity of recorded neurons by modelling activity as a linear noisy Gaussian observation *y*_*t*_ ∈ ℝ^*N*^ where *N* is the number of recorded neurons:

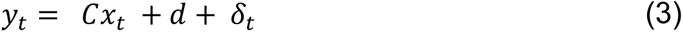

For *C* ∈ ℝ^*N*×*D*^ and *δ*_*t*_ ∼ *N*(0, *S*), a Gaussian noise term. Overall, the system parameters that rSLDS needs to learn consists of the state transition dynamics, library of linear dynamical system matrices and neuron-specific emission parameters, which we write as:

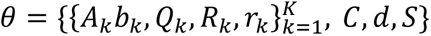

We evaluate model performance using both the evidence lower bound (ELBO) and the forward simulation accuracy (FSA) (Fig. 3a) described in Nair et al., 2023^15^ as well as by calculating the variance explained by the model on data.

We employed two-dimensional models, selecting the optimal number of states through 5-fold cross-validation. To ascertain which neurons contributed to each of the two model dimensions (x_1_ and x_2_), we initially confirmed the orthogonality of these dimensions by computing the subspace angle between them, (88.1 ± 0. 87°, n = 9 mice). Given this near orthogonality, we then utilized the columns of the emission matrix *C* to identify neurons that contributed to the two separate dimensions of the model.

#### Estimation of time constants

We estimated the time constant of each dimension of linear dynamical systems using eigenvalues *λ*_*a*_ of the dynamics matrix of that system, derived previously as^50^:

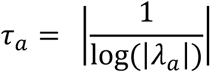

#### Calculation of line attractor score

To provide a quantitative measure of the presence of line attractor dynamics, we devised a line attractor score as defined in Nair et al., 2023 as:

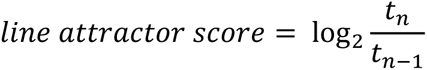

where *t_n_* is the largest time constant of the dynamics matrix of a dynamical system and *t_n_*_-1_ is the second largest time constant.

#### Calculation of auto-correlation half-width

We computed autocorrelation halfwidths by calculating the autocorrelation function for each neuron timeseries data (y_t_) for a set of lags as described previously^12^. Briefly, for a time series (y_t_), the autocorrelation for lag k is:

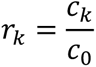

where c_*k*_ is defined as:

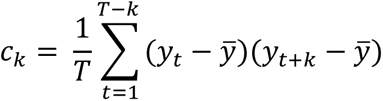

and c_0_ is the sample variance of the data.

#### Mechanistic modelling

We constructed a model population of N = 1,000 standard current-based leaky integrate-and-fire neurons as previously performed^22,36^. We first modelled a purely excitatory spiking network in which each neuron has membrane potential *x*_*i*_ characterized by dynamics:

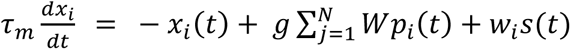

where *τ*_*m*_ = 20*ms* is the membrane time constant, *W* is the synaptic weight matrix, *s* is an input term representing external inputs and *p* represents recurrent inputs. To model spiking, we set a threshold (θ = 0.1), such that when the membrane potential *x*_*i*_(*t*) > θ, *x*_*i*_(*t*) is set to zero and the instantaneous spiking rate *r*_*i*_(*t*) is set to 1.

Spiking-evoked input was modelled as a synaptic current with dynamics:

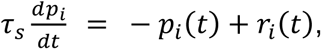

where *τ*_*s*_ is the synaptic conductance time constant. In excitatory networks, the network time constant *τ_n_* was derived as 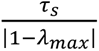, where *λ_max_* is the largest eigenvalue of the synaptic weight matrix *W*^34^.

We designed the synaptic connectivity matrix to include a subnetwork of 200 neurons (20% of the network), designated the integration subnetwork as suggested by empirical measurements, with varying densities of random connectivity as highlighted in Fig 3. Weights of the overall network were sampled from a uniform distribution: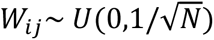, while weights of the subnetwork were sampled as 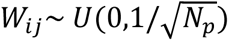, where *N*_*p*_ = 200.

External input was provided to the network as a smoothened step function consisting of four pulses at 20 ISI as provided in vivo. This stimulus drove a random 25% of neurons in the network.

To account for finite size effects and runaway excitation in networks, we also simulated models with fast feedback inhibition. This was modelled as recurrent inhibition from a single graded input *I*_*inh*_ representing an inhibitory population that receives equal input from and provides equal input to, all excitatory units. The dynamics of *I*_*inh*_ evolves as:

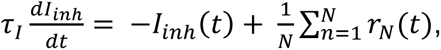

where *τ*_*I*_ = 50*ms* is the decay time constant for inhibitory currents. In this model, outside spiking events, the membrane potential evolved as:

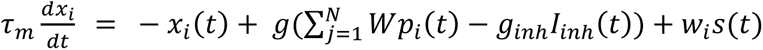

Model dynamics were simulated in discrete time using Euler’s method with a timestep of 1ms and a small gaussian noise term *η*_*i*_∼*N*(0,1)/5 was added at each time step. We used *g* = 1 and varied *g*_*inh*_ = 1,5,10 as suggested by measurements of inhibitory input to VMHvl^38^.

#### Statistical analysis

Data were processed and analyzed using Python, MATLAB, and GraphPad (GraphPad PRISM 9). All data were analyzed using two-tailed non-parametric tests. Mann-Whitney test were used for binary paired samples. Friedman test was used for non-binary paired samples. Kolmogorov-Smirnov test was used for non-paired samples. Multiple comparisons were corrected with Dunn’s multiple comparisons correction. Not significant (n.s), p > 0.05; *p < 0.05; **p < 0.01; ***p < 0.001; ****p < 0.0001.

## Acknowledgment

We thank Bin Yang for help in miniscope imaging and initial construct of the ChRminE plasmid, Itamar Landau for critical initial feedback, Tomomi Karigo for help in behavioral experiments, Jung-Sook Kim for help in stereotaxic surgeries and histology, Xiaolin Da for help in histology, Yi Huang for help with genotyping Maxi-prep, the Techlab at Caltech for help with 3D printing, Daniel Wagenaar for help with Hardware, Yonil Jung for help in first establishment of the 2P-SLM setup, Former Anderson lab members Brady Weissbourd, Ann Kennedy, Hui Chiu, Ke Ding, Kiichi Watanabe, Hidehiko Inagaki for initial feedback on this work, Current Anderson lab members for continuous feedback. Celine Chiu, Gina Mancuso, and Liliana Chavarria for lab management and administrative assistance. DJA is an investigator of the Howard Hughes Medical Institute. This work was supported by grants from the NIH (RO1MH112593, RO1MH123612 and RO1NS123916) and by the Simons Collaboration on the Global Brain. A.N is supported by a National Science Scholarship from the Agency of Science, Technology and Research, Singapore. A.V was supported by fellowships from the Human Frontiers Science Program and is a postdoctoral fellow at the Howard Hughes Medical Institute.

